# Koolen-de Vries Syndrome causal gene *KANSL1* is required for motile ciliogenesis

**DOI:** 10.1101/2024.11.14.621349

**Authors:** James D. Schmidt, Ananya Terala, Kate E. McCluskey, Belinda Wang, Anna C. Pfalzer, Helen Rankin Willsey

## Abstract

Koolen-de Vries Syndrome (KdVS), characterized by hypersociability, intellectual disability, and seizures, is caused by pathogenic variants in the gene *KANSL1*, which encodes a chromatin regulator in the NSL complex that also directly functions in mitotic spindle microtubule stability. Here we explored whether KANSL1 functions at the cilium, a microtubule-rich organelle critical for brain development, neuronal excitability, and sensory integration. Leveraging the *Xenopus* model, we found that Kansl1 is highly expressed in developing ciliated tissues and localizes within motile cilia. Moreover, *Kansl1* depletion caused ciliogenesis defects that could be partially rescued by human *KANSL1*. Based on these findings, we explored the prevalence of cilia-related clinical features including structural heart defects, hypogonadism, and structural respiratory defects among 99 individuals with KdVS, ages 1 month to 37 years old. To directly test if KdVS causes ciliary dysfunction in humans, we measured the well-established ciliary functional biomarker, nasal nitric oxide, in 11 affected individuals and observed a significant decrease when compared to unaffected family members. Together, this work establishes a ciliary contribution of *KANSL1* mutations to KdVS. This work adds to a growing literature highlighting the relevance of the cilium to neurodevelopmental disorders, particularly to those impacting sociability. Going forward, KANSL1 provides a unique opportunity to study a monogenic mechanism of hypersociability that may be useful in elaborating the molecular underpinnings of social behavior.

## INTRODUCTION

Koolen-de Vries Syndrome (KdVS) is a neurodevelopmental disorder characterized by hypersociability, dysmorphic facial features, epilepsy, intellectual disability, respiratory defects, kidney defects, congenital heart defects, hydrocephalus, and hypotonia (Koolen, Morgan, and de Vries 2023). KdVS is caused by pathogenic variants within the gene *KANSL1* (KAT8 Regulatory NSL Complex Subunit 1) or by microdeletions of its associated genomic locus 17q21.31 (Moreno-Igoa et al. 2015; Koolen, Morgan, and de Vries 2023). While KANSL1 is most well-characterized for its role as a chromatin regulator of KAT8 (Lysine Acetyltransferase 8), a histone acetyltransferase within the NSL (Nonspecific Lethal) Complex Subunit 1 (Mendjan et al. 2006), it is also known to regulate microtubule stability at the mitotic spindle (Meunier et al. 2015). Indeed, many other neurodevelopmental disorder-associated chromatin regulators (e.g., ADNP, HDAC6, SETD2, CHD3, and CHD4) share dual roles in regulating both histones and tubulins (Hubbert et al. 2002; I. Y. Park et al. 2016; Hadar et al. 2021; Oz et al. 2014; Sillibourne et al. 2007; Yokoyama et al. 2013; Simon et al. 2010). KANSL1 specifically has been shown to localize to centrosomes and bind directly to and stabilize microtubules at the mitotic spindle during mitosis, independent of its nuclear function (Meunier et al. 2015).

Precise regulation of microtubule stability is also critical for the complex process of cilia formation. Cilia are microtubule-based organelles present in almost all cells, including neurons, nucleated by modified centrosomes called basal bodies (Satir and Christensen 2007). Many of the clinical symptoms of KdVS are suggestive of altered ciliary function since cilia defects cause hydrocephalus, epilepsy, respiratory illnesses, heart defects, and kidney defects (Reiter and Leroux 2017). Therefore, we hypothesized that KANSL1 may also play a role in ciliary biology and that at least some of the KdVS clinical symptoms may be due to this function. Consistently, KANSL1 interacts with WDR5 (WD Repeat Domain 5), another member of the NSL complex (Sheikh, Guhathakurta, and Akhtar 2019; Mendjan et al. 2006), which is required for ciliogenesis independent of its histone function (Kulkarni et al. 2018). To test our hypothesis, we used *Xenopus*, a powerful *in vivo* model system for studying ciliary biology (Walentek 2021), to determine whether KANSL1 has a role at the cilium. Here we show that *kansl1* is expressed in highly ciliated tissues during embryonic development and localizes to ciliary axonemes and basal bodies of motile multiciliated cells of the epidermis. Loss of *kansl1* leads to motile cilia defects with a partial rescue following the introduction of human *KANSL1*. Consistent with a potential role for KANSL1 in ciliary biology, we document the prevalence of structural heart defects, structural lung defects, and hypogonadism within the KdVS affected population via caregiver surveys, confirming their presence. Finally, we assess nasal nitric oxide (nNO) levels, a biomarker for ciliary function, in individuals with KdVS and observe a significant decrease in nNO values compared to an unaffected family member. Together, these results establish that KANSL1 is involved in ciliary biology and that this function should be explored for therapeutic and diagnostic potential.

## RESULTS

### Kansl1 localizes to motile ciliary axonemes and basal bodies

Using whole-mount RNA *in situ* hybridization, we examined the expression pattern of *kansl1* in *X. tropicalis* during embryonic development. We observed strong *kansl1* expression in neural and ciliated tissues throughout development, particularly in the developing otic vesicle, pronephros (developing kidney), eye, and brain ventricles (**Fig. 1A**). *kansl1* was also highly expressed in the pharyngeal arches, neural crest-derived structures (**Fig. 1A**). Next, to assay subcellular protein localization, we tested two commercial Kansl1 antibodies by immunofluorescence staining of the multiciliated epidermis of *Xenopus laevis* and *Xenopus tropicalis* embryos. For both antibodies and in both species, we observed localization in the nucleus, as expected given its role in chromatin regulation (Mendjan et al. 2006). We also observed localization along ciliary axonemes labeled by Polyglutamylated or Acetylated α-Tubulin (**Fig. 1B-D, Fig. S1**), as well as at the base of cilia (basal bodies) labeled by Centrin-CFP (**Fig. 1E**). Substacked images show Kansl1 localization along ciliary axonemes labeled by Acetylated-α-Tubulin (**Fig. 1D**) and at basal bodies labeled by Centrin-CFP (**Fig. 1E**). To validate this localization, we depleted endogenous Kansl1 protein with a translation-blocking morpholino in *X. tropicalis* (marked by Centrin-CFP expression) and observed reduced ciliary staining compared to neighboring, unmanipulated (Centrin-CFP negative) cells (**Fig. S2**). We additionally validated this localization by expressing exogenous GFP-tagged human KANSL1 (hKANSL1-GFP) protein in this tissue in *X. laevis* and observed similar localization patterns along ciliary axonemes (**Fig. 1F**) and at ciliary basal bodies (**Fig. 1G**). Together, these results show that Kansl1 is highly expressed in developing embryonic ciliated tissues and localizes to ciliary structures in *Xenopus*.

**Figure 1.**
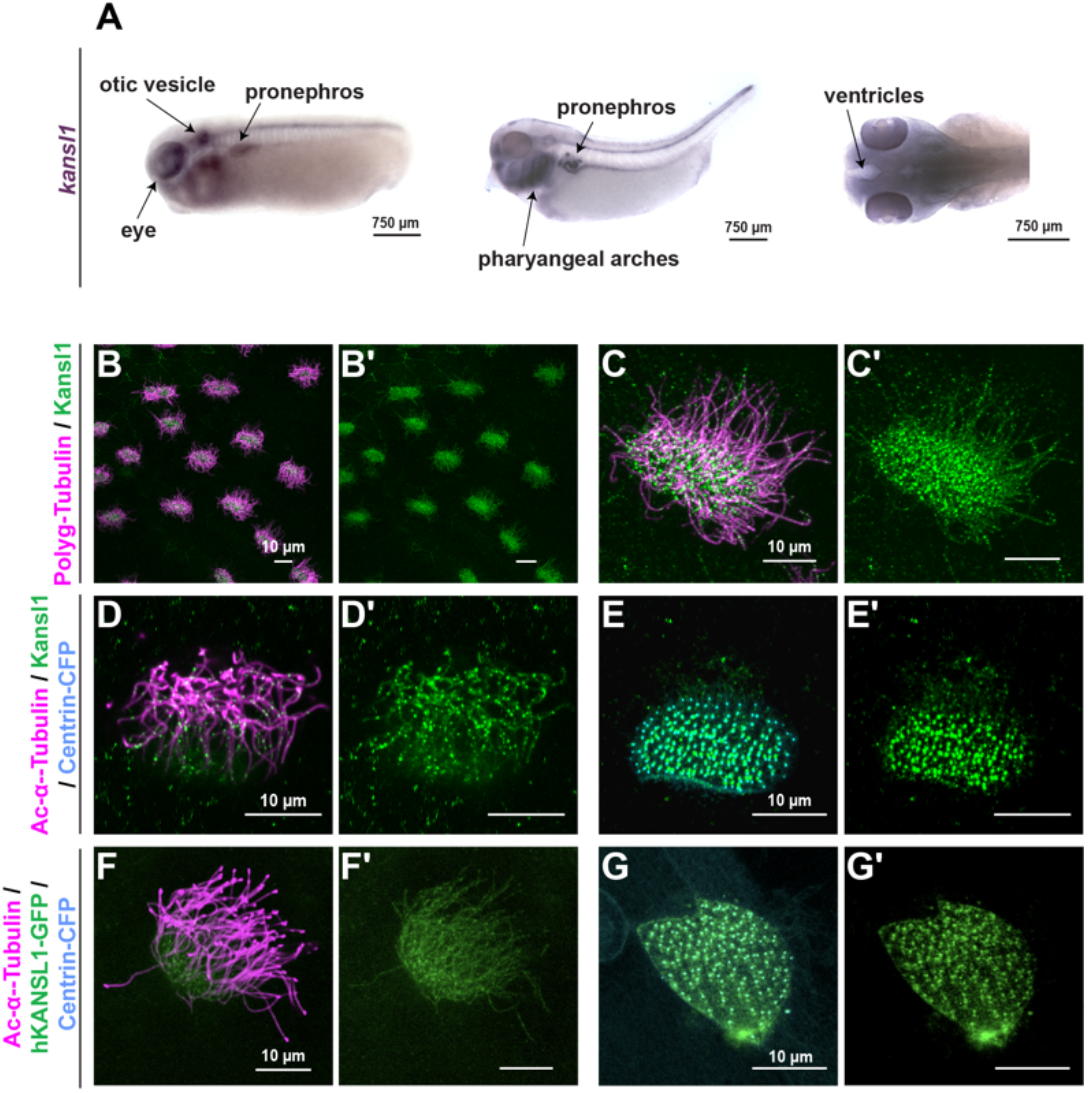
*kansl1* is expressed in ciliated tissues and localizes to cilia in *Xenopus* multiciliated cells. (A) *kansl1* is expressed in ciliated embryonic tissues including the pronephros, otic vesicle, eye, and brain ventricles in NF stages 25 and 30 *X. tropicalis* embryos. (B-B′) Kansl1 protein (green) localizes to cilia of *X. laevis* epidermal multiciliated cells labeled by Polyglutamylated-Tubulin (magenta) by antibody staining. (C-C′) Kansl1 (green) localizes as puncta along ciliary axonemes labeled by Polyglutamylated-Tubulin antibody staining (magenta). (D-D′) Kansl1 (green) also localizes to cilia labeled by Acetylated-α-Tubulin antibody staining (magenta, substacked to only show axonemal z-planes). (E-E′) Kansl1 (green) localizes to basal bodies labeled by Centrin-CFP (blue, substacked to only show basal body z-planes). (F-F′) Expression of human GFP-tagged KANSL1 (hKANSL1-GFP, green) localizes to ciliary axonemes labeled by Acetylated-α-Tubulin (magenta, substacked to only show axonemal z-planes). (G-G′) hKANSL1-GFP (green) also localizes to basal bodies labeled by Centrin-CFP (blue, substacked to only show basal body z-planes).

### Kansl1 is required for motile ciliogenesis

To investigate the developmental function of Kansl1 in ciliated cells, we used the same validated translation-blocking *kansl1* morpholino to deplete protein production in the *X. tropicalis* multiciliated epidermis and phenotyped cilia (**Fig. 2A-C**). To assay ciliary phenotypes, we quantified fluorescence intensity of the ciliary axonemal marker Acetylated-α-Tubulin of multiciliated cells and observed a statistically significant reduction following *kansl1* depletion that could be partially rescued by the co-injection of human KANSL1-GFP (**Fig. 2D**, both p < 0.0001 by one-way ANOVA and Dunnett’s test for multiple comparisons). We also observed a statistically significant reduction in apical area size following *kansl1* depletion that was also partially rescued by human KANSL1-GFP (**Fig. 2E**, both p < 0.0001 by one-way ANOVA and Dunnett’s test for multiple comparisons). These results show that Kansl1 is required for motile ciliogenesis in the *Xenopus* epidermis.

**Figure 2.**
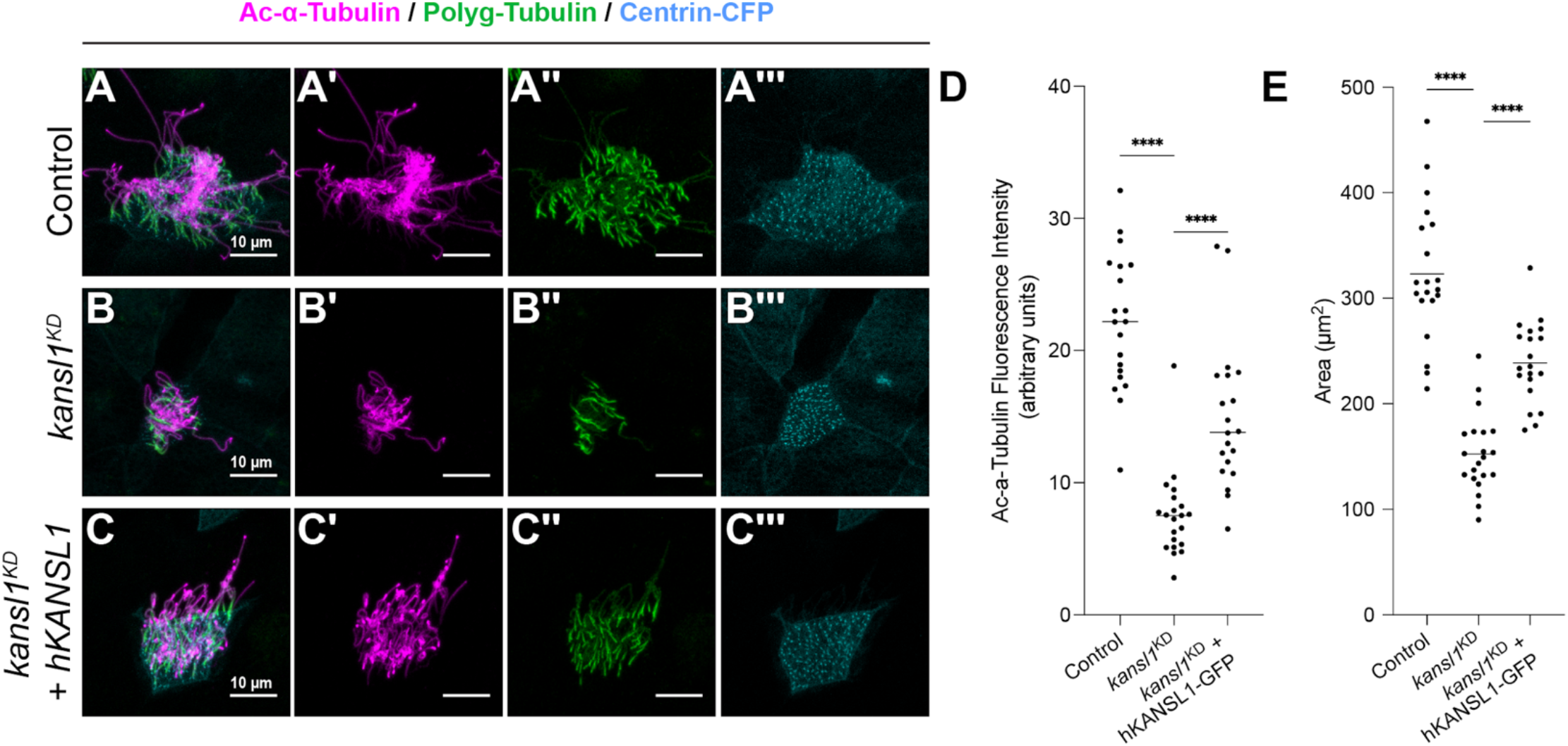
Depletion of *kansl1* results in motile ciliogenesis defects that can be partially rescued by human KANSL1. (A-C′′′) NF stage 26 *X. tropicalis* epidermis stained for Acetylated-α-Tubulin (magenta), Polyglutamylated-Tubulin (green), and concurrently expressing Centrin-CFP (blue). Depletion of *kansl1* by morpholino injection (B-B′′′) results in motile ciliogenesis phenotypes when compared to a control morpholino (A-A′′′). This effect can be partially rescued by co-injecting a human GFP-tagged KANSL1 (hKANSL1-GFP) (C-C′′′). (D) Quantification of Acetylated-α-Tubulin fluorescence intensity indicates that depletion of *kansl1* causes a significant decrease in signal compared to control conditions, which can be partially rescued by supplying hKANSL1-GFP. (E) Quantification of apical area size indicates that depletion of *kansl1* causes a significant decrease compared to control injections, which can be partially rescued by supplying hKANSL1-GFP. ****p < 0.0001 by a one-way ANOVA followed by a Dunnett’s test for multiple comparisons.

### Prevalence of motile ciliopathies within the KdVS affected population

KdVS has been associated with a range of clinical presentations consistent with ciliary dysfunction, including distinctive facial features, seizures/epilepsy, congenital heart defects, urogenital anomalies, and musculoskeletal anomalies (Koolen, Morgan, and de Vries 2023). However, these observations have been limited to broad clinical categories with general estimates of prevalence (e.g., “very common” = over 75%, versus “common” = 50-75%) (Koolen, Morgan, and de Vries 2023). Therefore, we wanted to also probe within these general categories to determine the prevalence of specific ciliary-related clinical presentations. To confirm these observations and add more granularity to the available phenotypic literature, we analyzed available caregiver survey data collected by COMBINEDBrain reporting on 99 affected individuals ranging in age from 1 month to 37 years old, with 48% female and 52% male. We observed high rates of multi-organ findings, including vision (45%), skeletal (80%), urogenital (41%), respiratory (60%), and cardiac abnormalities (44%), as well as seizures (47%) (**Fig. 3A**). These prevalences are in range with what is currently published (Koolen, Morgan, and de Vries 2023), except that we observe higher rates of skeletal defects most likely because we asked targeted questions about it.

**Figure 3.**
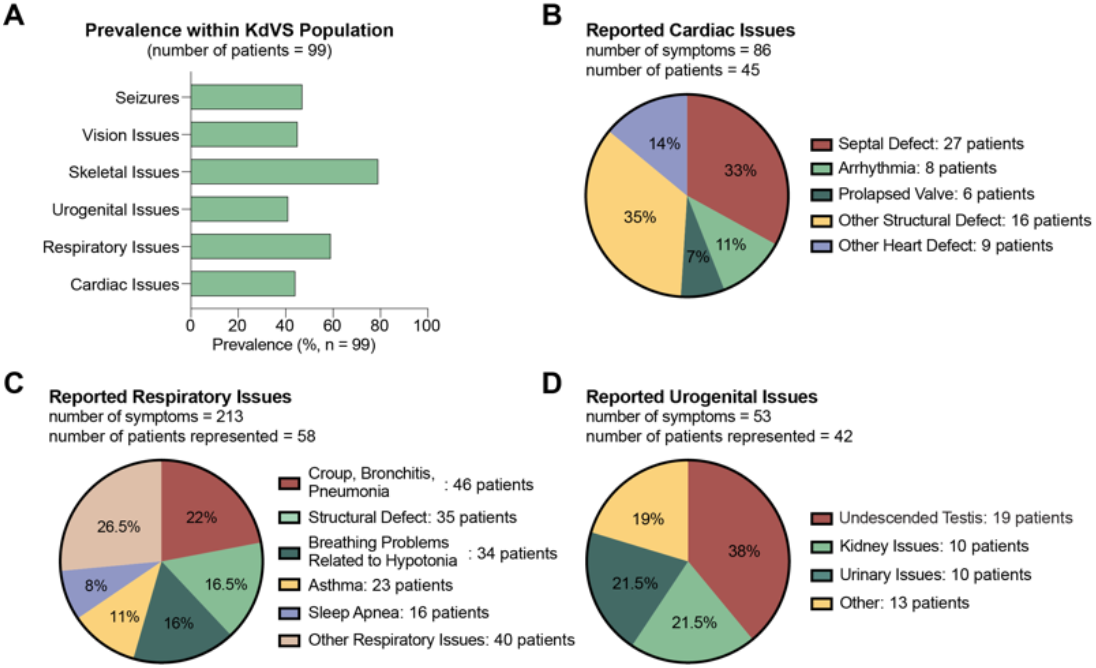
Prevalence of clinical features among 99 individuals with KdVS. (A) Caregiver surveys representing 99 individuals with KdVS reveal prevalence of clinical symptoms affecting many organ systems, consistent with a potential ciliary contribution to pathogenesis. (B) Of the 86 reported cardiac symptoms from 45 affected KdVS individuals, 75% were structural abnormalities (septal defect, prolapsed valve, and other), 11% were arrhythmias, and 14% were other heart defects. (C) Of the 213 reported respiratory symptoms from 58 KdVS individuals, 35% were breathing problems (related to hypotonia, asthma, and sleep apnea), 22% were related to infections, 16.5% were structural defects and 26.5% were listed as other respiratory issues. (D) Of the 53 reported urogenital symptoms from 42 KdVS individuals, 38% were undescended testis, 21.5% were kidney issues, 21.5% were urinary issues, and 19% were listed as other. See Table S1.

To characterize more specific conditions within these categories directly linked to ciliary issues, we probed the prevalence of structural heart defects, structural lung defects, and hypogonadism (Djenoune et al. 2022; Kim et al. 2018; Boon et al. 2015). Out of all the symptoms reported for cardiac issues, 75% were structural heart defects (**Fig. 3B, Table S1**), representing 45% of the assessed KdVS population. Within respiratory issues, structural respiratory tract defects amounted to 16.5% (35% of the population) and respiratory infections were 22% (47% of the population) (**Fig. 3C, Table S1**). Within urogenital issues, undescended testis, a feature of hypogonadism, were 38% of the reported urogenital symptoms (19% of the population, 37% of males) (**Fig. 3D, Table S1**) and kidney issues accounted for 21.5% of the reported symptoms (10% of the population) (**Fig. 3D, Table S1**). Ciliopathies are well-established to lead to structural heart defects, structural lung defects, and hypogonadism (Reiter and Leroux 2017). These rates of reported symptoms with potential underlying ciliary contributions is consistent with our hypothesis that KANSL1 functions in ciliary biology and that this function is relevant to KdVS.

### Individuals with KdVS have significantly lower nasal nitric oxide than unaffected family members

Our data confirms that a substantial portion (at least 35%) of KdVS individuals exhibit phenotypes consistent with ciliopathies. Consequently, we sought to test whether individuals diagnosed with KdVS show disrupted biomarkers of ciliary function. One common biomarker for human ciliary function, used to diagnose the ciliopathy primary ciliary dyskinesia, is nasal nitric oxide (nNO) (Leigh et al. 2013; Beydon et al. 2023). nNO is produced by ciliary beating from multiciliated cells of the nasal epithelium, a cell type in which *KANSL1* is expressed in humans (**Fig. 4A**). In ciliopathies, cilia formation and/or beating is disrupted, leading to reduced nNO production (**Fig. 4B**) (Wodehouse et al. 2003; Narang et al. 2002). Therefore, we measured nNO production in 11 individuals with KdVS (ages 2 to 30), compared to an unaffected immediate family member to control for genetic background (ages 10 to 63). Individuals with KdVS had significantly lower nNO values compared to their family members (**Fig. 4C**, p = 0.0013 by parametric one-tailed t test). The effect was also statistically significant when tested unpaired (p = 0.0051 by parametric one-tailed t test). The nNO differences observed here were not due to differences in age or sex between tested individuals (**Figs. 4D, S3**). Together, our results demonstrate that KANSL1 is required for ciliogenesis in *Xenopus* and that human individuals with KdVS have clinical symptoms consistent with a ciliary contribution to pathobiology and reduced nNO levels, a biomarker of motile cilia function.

**Figure 4.**
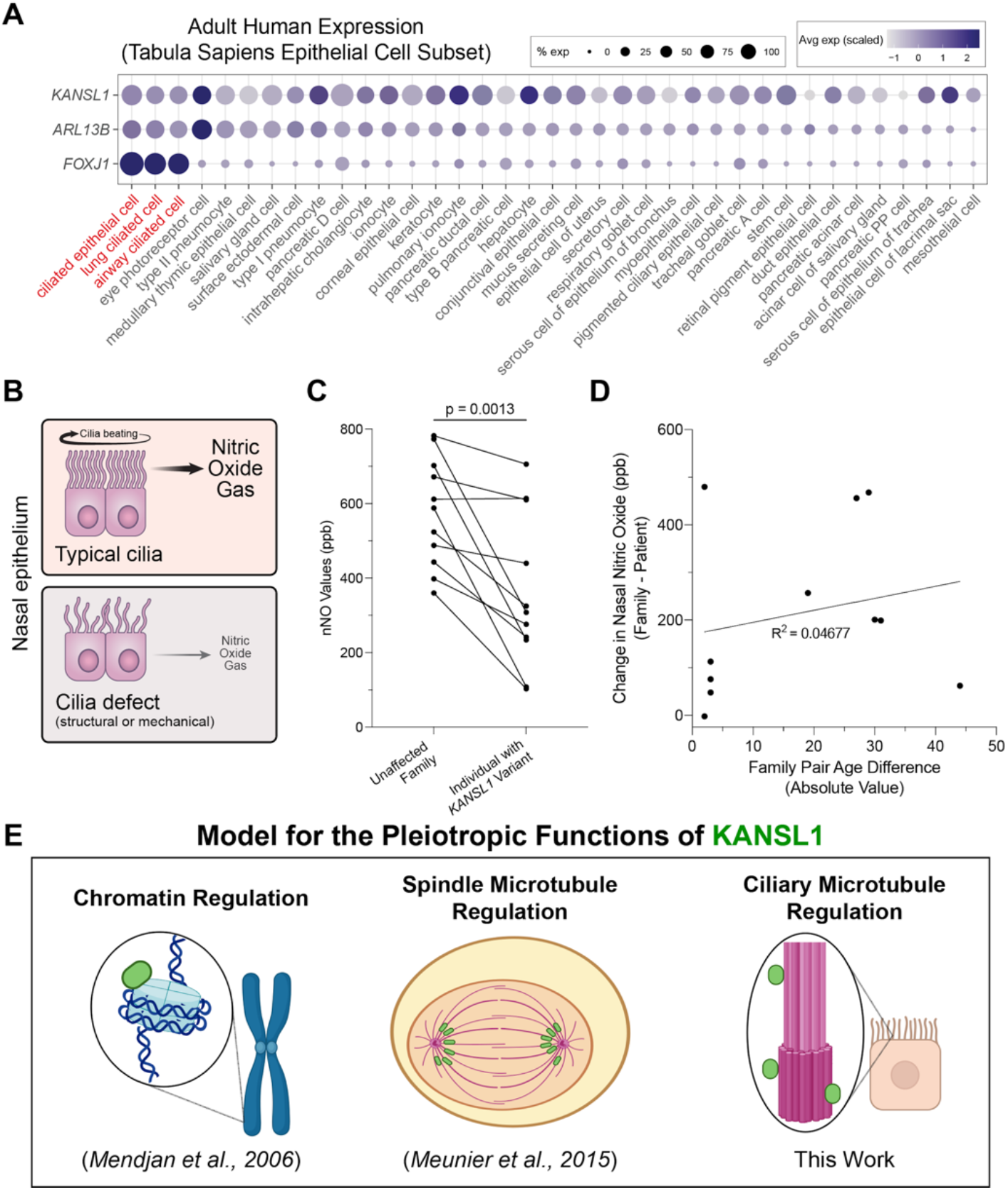
Ciliary biomarker disruption in KdVS supports a new model for KANSL1 functional pleiotropy. (A) *KANSL1* expression in relation to two ciliary marker genes, *ARL13B* and *FOXJ1*, in epithelial cells from human Tabula Sapiens Dataset with expression in airway ciliated cells (Tabula Sapiens Consortium* et al. 2022). Dot size represents percent of cells expressing the gene of interest while purple intensity represents scaled average expression. *KANSL1* is expressed in airway ciliated cells marked by high *FOXJ1* expression. (B) Schematic illustrating that ciliary beating produces nitric oxide gas in the nasal epithelium of humans, which is disrupted in the context of cilia defects. (C) nNO values are significantly lower in individuals with KdVS compared to an unaffected family member, p = 0.0013 (parametric one-tailed t test). (D) Plot of change in nNO vs. Age difference between KdVS individuals and their family members demonstrates minimal linear correlation, R^2^ = 0.04677. (E) Previous work has shown KANSL1 to function as a chromatin regulator at histones and a microtubule stabilizer at the mitotic spindle (Mendjan et al. 2006; Meunier et al. 2015). This work adds a new function of KANSL1 in regulating ciliary biology.

## DISCUSSION

Given the documented role of KANSL1 as a chromatin regulator (Mendjan et al. 2006), conventional wisdom has suggested that its mechanism of action with respect to KdVS lies in regulating gene expression. However, KANSL1 also directly binds and stabilizes microtubules (Meunier et al. 2015), the main structural component of cilia. Consistently, here we show that KANSL1 localizes to and functions at motile cilia in the *Xenopus* epidermis and that individuals with variants in this gene have a reduced ciliary functional biomarker. Therefore, here we offer the additional possibility that the role of KANSL1 in ciliary biology also contributes to the clinical presentation of KdVS. We propose a model in which KANSL1 functions as a chromatin regulator in the nucleus, while in the cytoplasm and the cilium it functions to regulate microtubule stability (**Fig. 4E**). The relative contributions of each function to KdVS is unclear and parsing these functions will be important future work that may be informed by investigating patient-derived missense variants in *KANSL1* that have function-specific effects, if they exist, as has been done previously with *WDR5* (Kulkarni et al. 2018). This dual function in histone and tubulin regulation is also well-established for many other genes associated with neurodevelopmental disorders including *SETD2, ADNP, HDAC6, CHD3*, and *CHD4* (Hubbert et al. 2002; I. Y. Park et al. 2016; Hadar et al. 2021; Oz et al. 2014; Sillibourne et al. 2007; Yokoyama et al. 2013; Simon et al. 2010), suggesting that other such proteins may also have roles at the cilium that have yet to be described. Consistently, we have observed many chromatin regulators associated with autism spectrum disorder localizing to microtubules of the mitotic spindle (Lasser et al. 2023). The work shown here adds to a growing literature linking rare neurodevelopmental disorder-associated genes with tubulin and ciliary biology, motivating continued work investigating potential ciliary functions of such genes (Willsey et al. 2020, 2018; Teerikorpi et al. 2024; Sun et al. 2024; Marley and von Zastrow 2010).

Of special interest is the high prevalence of hypersociability documented among individuals with KdVS (Toth 2019). Identifying the molecular underpinnings of sociability is important, and significant progress has been made in characterizing rare mutations associated with decreased sociability and other neurodevelopmental disorders (Willsey et al. 2022), but the genetics and biology of increased sociability is far less well understood. Williams Syndrome, another developmental disorder characterized by hypersociability, has been the paradigmatic example that demonstrated that social reciprocity and intellectual ability were separable (Beuren et al. 1964; Jones and Smith 1975; Vonarnim and Engel 1964). KANSL1 supports that conclusion and provides some unique opportunities to delve further into social functioning at the molecular level given the challenges of a contiguous gene syndrome like Williams Syndrome (Crespi and Procyshyn 2017; Zhou et al. 2022). In contrast, KdVS can be caused by a single variant within *KANSL1*, making this gene particularly attractive for studying the molecular mechanisms of sociability. Consistently, some murine models of KdVS have shown hypersociability behaviors (Arbogast et al. 2017), suggesting that animal models may be useful in this effort. The ciliary function identified for KANSL1 here also adds to a growing body of literature converging on the cilium as a relevant organelle for genes associated with atypical sociability (Marley and von Zastrow 2010; Teerikorpi et al. 2024; Willsey et al. 2020, 2018). Given the high prevalence of autism among individuals with pathogenic variants in ciliary genes (Ozonoff et al. 1999; Kerr, Bhan, and Héon 2015; Sacai et al. 2020), future work dissecting the role of cilia in social behavior may be illuminating.

While this paper establishes the function of KANSL1 in ciliary biology, extensive future work is required to dissect this role and translate its relevance to clinical practice. Nevertheless, despite our functional work being done in *Xenopus*, we were able to make predictions that were validated in a human cohort by nNO testing. While nNO is used as a diagnostic biomarker for the ciliopathy primary ciliary dyskinesia (Beydon et al. 2023), this is the first study to propose its use for identifying new genetic ciliopathies. Using this noninvasive, 30 second test we observed encouraging results that this metric may be valuable in the context of identifying genetic ciliopathies and potentially in providing a quantitative value for assessing future therapeutic efficacy. Of note is that only 1 individual with KdVS had a value close to the diagnostic threshold for primary ciliary dyskinesia (103 ppb, when the diagnostic threshold on this machine is < 100 ppb (Paternò et al. 2023; Beydon et al. 2023)), which is unsurprising given the heterozygous, *de novo* nature of the variants of this genetic disorder, compared to primary ciliary dyskinesia which usually results from inherited, homozygous loss of function variants (Horani and Ferkol 2021). Nevertheless, more work is required to determine the stability of these values over time within this cohort and any potential interactions with medication status.

Here we have demonstrated that KANSL1 is involved in motile multiciliogenesis in the *Xenopus* epidermis, but cilia exist in myriad tissues and change over development with a correspondingly wide array of cellular functions (Hansen et al. 2024; Drummond 2012). Of relevance to KdVS are likely: motile multiciliated cells of brain ependymal glia given the prevalence of hydrocephalus; motile multiciliated cells of the respiratory tract given the prevalence of bronchiectasis and reduced nNO; motile monociliated cells of the left-right embryonic organizer given the prevalence of structural heart defects; nonmotile primary cilia in neurons given the prevalence of seizures; and primary cilia of the migrating neural crest given the prevalence of dysmorphic facial features. All of these contexts will be important future areas of investigation. Nevertheless, the insight that KANSL1 is involved in ciliary biology has immediate implications for affected individuals, including the likelihood of underdiagnosis of kidney defects (as previously seen in DYRK1A Syndrome (Blackburn et al. 2019)) and the potential for affected individuals to be seen by cilia centers that may be able to provide more tailored care. This work also predicts potential therapeutics and provides a screenable phenotype for future drug repurposing efforts. Of potential interest are microtubule stabilizing drugs which could be repurposed from oncology as well as histone deacetylase (HDAC) inhibitors, particularly those that affect HDAC6, since HDAC6 is known to regulate ciliary acetylation of tubulin, a stabilizing post-translational modification (Yoon and Eom 2016; Hubbert et al. 2002; Ran et al. 2015; Mergen et al. 2013). Recent work has also identified HDAC inhibitors as potential therapeutics in other neurodevelopmental disorders, but with the assumed mechanism of action being epigenetic (Hull, Montgomery, and Leyva 2016; Bose, Dai, and Grant 2014).

In sum, the work presented here establishes that KANSL1 is involved in motile ciliary biology and that this function may be relevant to the clinical presentations observed in KdVS. Given the hypersociability of individuals with variants in *KANSL1*, this gene may be particularly informative in identifying the molecular mechanisms underlying social behavior, with the possibility that ciliary biology is relevant. As we move forward, parsing the pleiotropic and potentially cell-type-specific roles of KANSL1 in the nucleus and at microtubule-based structures like the cilium will be critical toward this effort as well as for alleviating clinical symptoms in KdVS.

## MATERIALS AND METHODS

### *Xenopus* husbandry and animal care

Both *Xenopus laevis* and *Xenopus tropicalis* were used in this study. *In situ* hybridization, morpholino, and rescue experiments were done in *X. tropicalis*. Protein localizations were performed in *X. laevis* and *X. tropicalis*. All frog care was performed according to UCSF IACUC protocol #AN199587-00A. All *Xenopus* stages were based off of (Nieuwkoop and Faber 1956). Xenbase (RRID:SCR_003280) was used for anatomical resources, phenotypes, and genetic references (M. Fisher et al. 2023; M. E. Fisher et al. 2022; Segerdell et al. 2013). Wildtype frogs were supplied by the National Xenopus Resource Center (RRID: SCR_013731) (Pearl et al. 2012).

### *Xenopus* whole mount RNA *in situ* hybridization

*X. tropicalis kansl1* probe plasmid (XGC Clone 7691711) was a kind gift from Dr. Richard Harland (UC Berkeley). Digoxigenin-11-UTP-labeled antisense RNA probe was synthesized from this plasmid by standard protocol (Sive 2000) using SalI restriction enzyme and T7 polymerase. Whole mount RNA *in situ* hybridization was performed on *X. tropicalis* stage 25 and 30 embryos according to (Willsey 2021) with the omission of the proteinase K step.

### *Xenopus* immunofluorescence

The following primary antibodies were used: Acetylated-α-Tubulin (Sigma T6793, 1:500), Polyglutamylated-Tubulin (AdipoGen Life Sciences AG-20B-0020-C100, 1:500), KANSL1 (Abcam ab230008, 1:250 and Abnova PAB20355, 1:250). The following secondary antibodies were used at 1:250: anti-mouse-555 (Thermo Fisher Scientific, A32727), anti-mouse-647 (Thermo Fisher Scientific, A32728), anti-rabbit-555 (Thermo Fisher Scientific, A32732), and anti-rabbit-647 (Thermo Fisher Scientific, A32733). All samples were fixed in 4% paraformaldehyde for 45 minutes then underwent standard immunofluorescence staining (Willsey 2021) with the omission of bleaching. Phalloidin (Life Technologies A22287, 1:250) and DAPI (1:1000) were incubated along with the secondary antibodies.

### Human protein localizations in *Xenopus*

hKANSL1-GFP plasmid was constructed by cloning NM_001193466 cDNA into pcDNA3.1+ at the N-terminal cloning site. 20 pg of hKANSL-GFP plasmid was injected into 1 blastomere of the 2-cell stage along with 100 pg of Centrin-CFP mRNA (T. J. Park et al. 2008; Antoniades, Stylianou, and Skourides 2014). Multiple concentrations of hKANSL1-GFP plasmid were tested, with 20 pg per blastomere being the lowest amount visible without causing overexpression phenotypes. hKANSL1-GFP was also injected alone to ensure that the localizations were not due to the presence of Centrin-CFP or due to spectral overlap, which they were not.

### *Xenopus* gene perturbations and rescues

For *Xenopus* knockdown experiments, morpholinos were purchased from Gene Tools, and their sequences are: *kansl1* (5’-GAGCCATCGCAGCCATTCAG-3’) and standard control (5’-CCTCTTACCTCAGTTACAATTTATA-3’). For both *kansl1* and the standard control morpholino, 3.32 ng of morpholino was injected into 1 blastomere of 2-cell stage *X. tropicalis* embryos, along with 100 pg of tracer Centrin-CFP mRNA. Rescue conditions were injected with 3.32 ng of morpholino with 40 pg of hKANSL1-GFP into 1 blastomere of 2-cell stage *X. tropicalis* embryos.

### Imaging and image analysis

All images were acquired on a Zeiss LSM 980 confocal microscope with 63x oil objective. Images were acquired in confocal mode. Images were processed in ImageJ (v2.0.0). Positively-injected ciliated cells were identified by positive Centrin-CFP signal. Acetylated-α-Tubulin and Polyglutamylated-Tubulin fluorescence intensity were quantified in ImageJ (v2.0.0) with the measure tool. At least 3 representative embryos per condition were assessed, with at least 20 cells per condition. Apical area was manually measured in ImageJ (v2.0.0) using the measure tool. Statistical analyses to compare the morpholino and rescue conditions to the control consisted of testing for normality via the Anderson-Darling test, followed by a one-way ANOVA and a Dunnett’s test for multiple comparisons. All statistical tests were performed in Prism (v10.2.0).

### KdVS caregiver surveys

The Koolen-de Vries Syndrome Foundation designed an online survey to better characterize select symptoms not previously associated with KdVS. The 28-question survey was completed by KdVS community members at the bi-annual KdVS Scientific and Patient Advocacy Summit in Orlando, Florida, between July 19th-21st, 2023 and subsequently distributed through various online KdVS social media platforms using the REDCap electronic data capture tools hosted at COMBINEDBrain (Harris et al. 2019). The survey included questions related to the presence of epilepsy, cardiac, respiratory, skeletal, vision and urogenital abnormalities using yes/no questions. Caregivers endorsing particular symptoms were then presented with a table of detailed descriptions for that symptom to provide likert-scale responses: previously had, currently having, never had, not sure. Data shown here includes the overall percentage of caregivers who endorsed select symptoms. The study was approved by the NorthStar Review Board (#NB200058). Consent was determined to be voluntary completion of the study survey. All data included in this study were analyzed in a de-identified manner.

### Expression of *KANSL1* versus ciliated cell markers across cell types

Tabula Sapiens (human cell atlas) dataset: We downloaded the Tabula Sapiens 2022 (Tabula Sapiens Consortium* et al. 2022) epithelial subset single-cell RNAseq Seurat object from CELLxGENE (https://cellxgene.cziscience.com/e/97a17473-e2b1-4f31-a544-44a60773e2dd.cxg/) (Megill et al. 2021) in June 2023. This dataset contains scRNAseq data for 104,148 cells across 18 tissues and 62 cell types, 3 of which are explicitly classified as ciliated (ciliated epithelial cell, lung ciliated cell, ciliated cell). Gene expression levels (UMI) were previously quantile normalized. We grouped cells by “cell_type”, and used percent expression (pct.exp, percent of cells in each cell type that have detected expression of the gene of interest) as a proxy for gene expression level. We categorized cell types into specific tissues of origin based on the “tissue_in_publication” that contributes the largest proportion of cells to each cell type. We renamed “ciliated cell” to “airway ciliated cell” as all the cells from this cell type derived from the trachea (**Fig. 4A**). We show data only from the 8 tissues (with 37 associated cell types) that had at least one cell type with above 20% expression of *FOXJ1* or *ARL13B*.

### nNO measurements and analyses

This work was conducted under the advisement of the UCSF Institutional Review Board with approval #23-39385. A caregiver provided age, genetic diagnosis, and sex. KdVS individual and unaffected family member (sibling or parent) values were recorded using a NIOX VERO nNO machine by tidal breathing method using adult or pediatric olives, as appropriate. Tidal breathing method, which requires minimal cooperation compared to the forced exhale method, was selected since many of these diagnosed individuals have intellectual disability. All study participants were able to complete the measurement. This machine has been shown to discriminate primary ciliary dyskinesia from controls with 100% sensitivity and specificity (Jõgi et al. 2023). For the tidal breathing method we used, a cutoff value of 30 nL/min (< 100 ppb on this machine) has been proposed for diagnosing primary ciliary dyskinesia (Beydon et al. 2023; Paternò et al. 2023). One diagnosed individual was close to this value with a measurement of 103 ppb. Statistical differences in nNO between groups were assessed by parametric, one-tailed, paired and unpaired Wilcoxon tests. We went into this work with the hypothesis that KdVS individuals would have lower nNO values, justifying a one-tailed test. Statistical differences in change of nNO between same sex and opposite sex pairing for control was assessed via a nonparametric, two-tailed, unpaired Mann-Whitney test. All statistical tests were performed in Prism (v10.2.0).

## ACKNOWLEDGEMENTS

We thank: the KdVS community for participating in this research; Ashley Point and the KdVS Foundation for insight and advice; Matthew State for invaluable feedback; Nolan Wong and UCSF LARC for animal care; Ethel Bader, Catherine Nguyen, Juan Arbelaez, Jean Dea, and Milagritos Alva for laboratory maintenance and support; Ashley Clement, Gigi Paras, Sonia Lopez, and Linda Chow for administrative support; Christina Roca and Micaela Lasser for preliminary localizations of KANSL1 in *Xenopus*; and Ngoc Ly for suggesting nasal nitric oxide as a potentially useful ciliary biomarker. This work would not be possible without daily reference to the *Xenopus* community resource Xenbase (RRID:SCR_003280) and expertise and frog resources from the National *Xenopus* Resource (RRID:SCR_013731).

## AUTHOR CONTRIBUTIONS

Conceptualization: HRW

Formal Analysis: JDS

AT Funding Acquisition: HRW

ACP Investigation: JDS, KEM, AT, ACP, BW

Methodology: JDS, HRW, AT, ACP

Project Administration: HRW

Resources: HRW, ACP, AT

Supervision: HRW, ACP

Validation: JDS

Visualization: JDS, KEM, HRW

Writing-original draft: JDS

Writing-review & editing: JDS, HRW, KEM, ACP, BW

## DATA AND MATERIALS AVAILABILITY

All data are within this manuscript.

## SUPPLEMENTAL FIGURES

**Figure S1.**
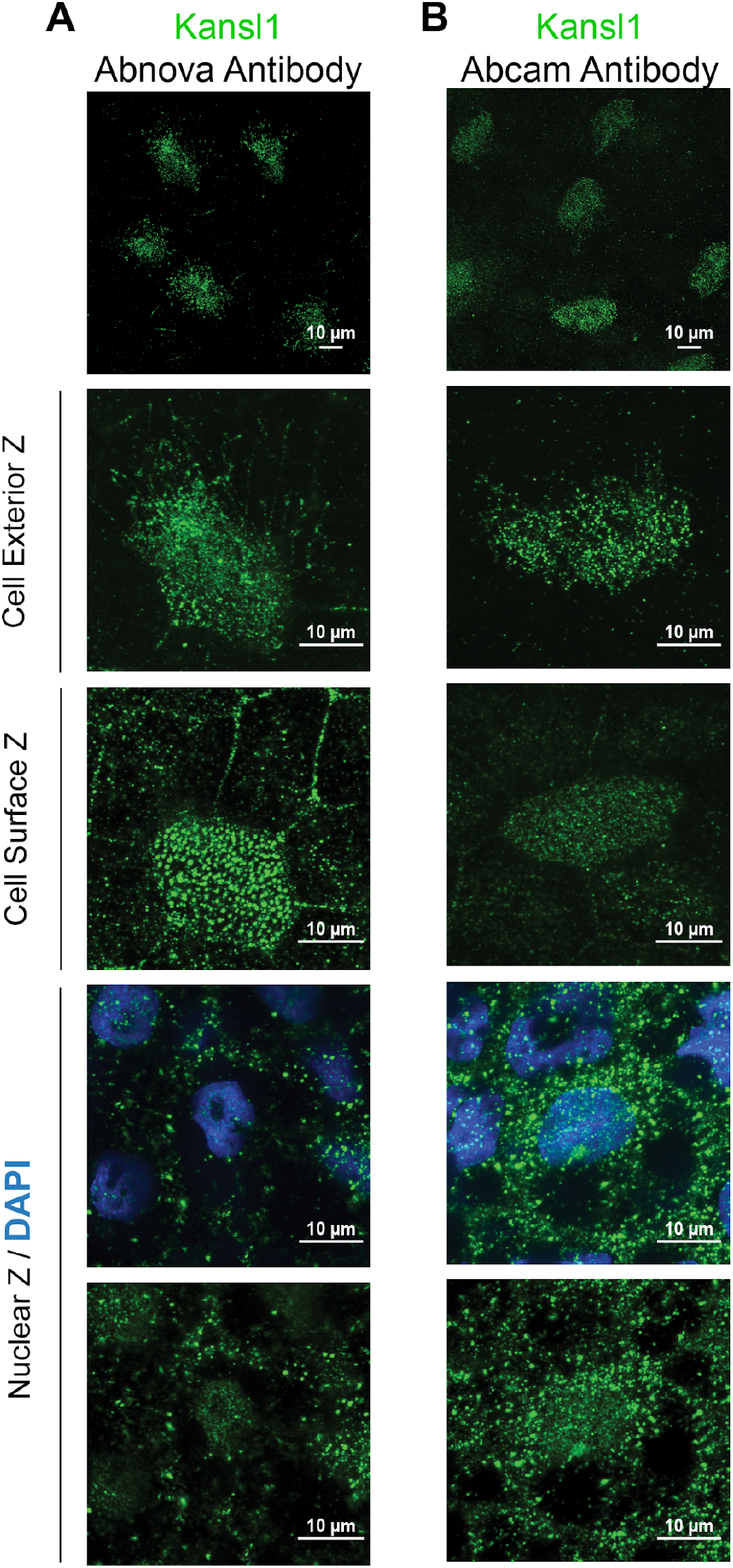
Two independent commercial Kansl1 antibodies support the localization of Kansl1 in the nucleus and on cilia in *Xenopus*. (A) Abnova Kansl1 antibody (green) and DAPI (blue) shown on cell exterior, cell surface, and nuclear level. (B) Abcam Kansl1 antibody (green) and DAPI (blue) shown on cell exterior, cell surface, and nuclear level. These stainings are from *X. laevis*, but similar results were observed in *X. tropicalis* for both antibodies. Stainings were done in the presence or absence of ciliary co-markers (Acetylated-α-Tubulin or Centrin-CFP) and both experiments showed similar results, supporting these localizations.

**Figure S2.**
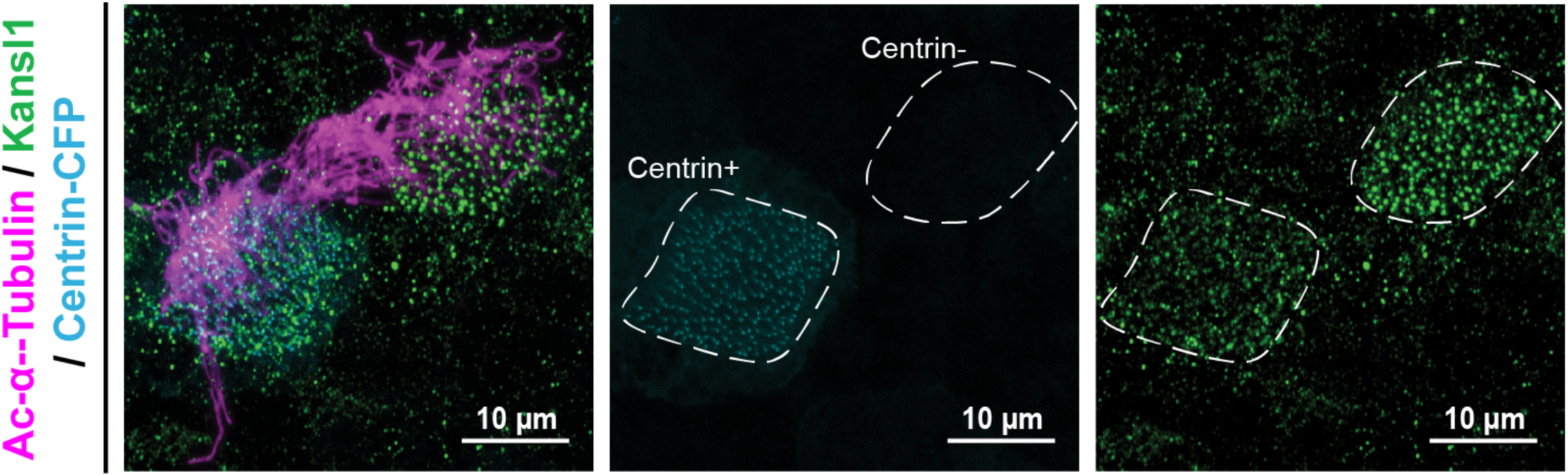
*kansl1* depletion reduces Kansl1 antibody staining. *X. tropicalis* epidermal multiciliated cells co-injected with *kansl1* morpholino and Centrin-CFP (blue) show a decrease in Kansl1 antibody staining (green, Abnova antibody) when compared to an adjacent unmanipulated cell (Centrin-negative), supporting the fidelity of both reagents.

**Figure S3.**
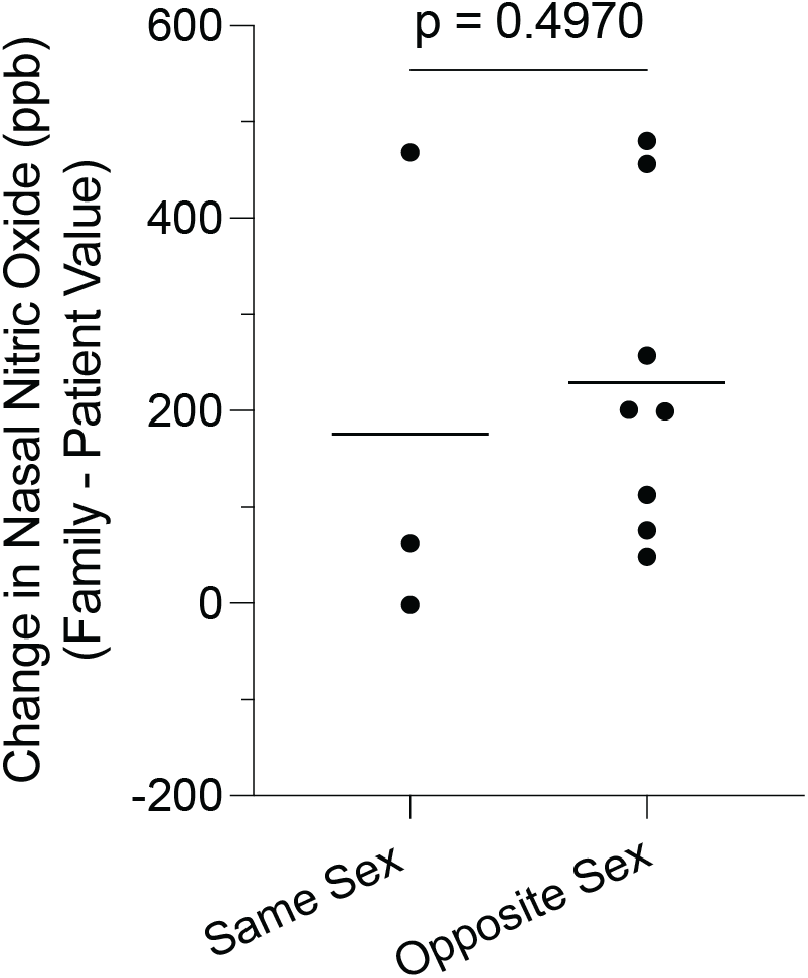
nNO differences are independent of sex. Differences in sex matching between family members does not affect nNO results (p = 0.50, unpaired, nonparametric Mann-Whitney rank sum test).

**Table S1. Breakdown of clinical features from KdVS caregiver survey data**. Tabs include specific breakdown data for cardiac, respiratory, and genitourinary responses.

